# Grain filling leads to backflow of surplus water in maize grain via the xylem to the cob and plant

**DOI:** 10.1101/2022.07.22.501090

**Authors:** Gui-Ping Zhang, Mukti Marasini, Wei-Wei Li, Feng-Lu Zhang

## Abstract

The rapid dehydration rate of maize grain is one of the main characteristics of cultivar selection of mechanical grain harvest, but the dominant driving force and mechanism of grain dehydration before physiological maturity remains disputable and obscure, respectively. This study found that, from grain formation to 5-10 days before physiological maturity of early and middle maturity maize cultivars, the main driving force of grain dehydration is filling and then converts to surface evaporation, by comparing the grain moisture content and dehydration rate between grain coated treatment and control. In the dye movement experiment, xylem-mobile dye movement into grain through pedicel xylem was observed during grain formation period, and declined and gradually not observed after grain formation. Xylem-mobile dye movement out of ear via cob, ear-pedicel and stem xylem was observed after grain formation. In addition, from grain formation to physiological maturity, there was a very significant positive correlation between grain filling rate and dehydration rate. According to these results, it is proposed here that in the grain dehydration phase driven by filling, the surplus water in grain flows back to cob via pedicel xylem, and some of it flows back to plant via cob and ear-pedicel xylem.

**Highlight:** The surplus water in grain driven by grain filling flows back to the cob and plant for recyclingvia the xylem during the development of maize grain.

## Introduction

The maize*(Zea mays L.)* grain mechanical harvesting system combines ear picking and threshing at one time, which reduces the operation link and the labor input, and improves the production efficiency (Guo *et al*., 2017). The maize grain mechanical harvest is the development direction of maize harvest (Ministry of Agriculture and Rural Affairs Information Office, 2021), as well as the key to understand the whole mechanization of maize and change the mode of production. At present, the high breakage rate of grains is one of the main problems restricting the popularization and application of maize grain mechanical harvesting technology (Chai *et al*., 2017), and the grain moisture content at harvest is the main factor affecting the breakage rate (Wang *et al*., 2017; Guo *et al*., 2022; Shi *et al*., 2022). The maize grain moisture content at harvest in China is universally on the high side(≥ 30%), which is not suitable for mechanical grain harvesting (Li *et al*., 2021a). Dehydration is one of the key component of the processes of grain ripening and drying, which determines the grain moisture content at harvest (Kang *et al*., 1986; Muchow *et al*., 1990). Currently, there are numbers of studies on the influencing factors of grain dehydration, such as meteorological factors, variety properties and cultivation measures (Zhang *et al*., 2018; Gao *et al*., 2018; Zhao *et al*., 2018; Wang *et al*., 2019), but few studies focused on the mechanism of grain dehydration. It has been argued that grain dehydration is related to grain filling (Kang *et al*., 1986; Cross *et al*., 1991; Li *et al*., 2014; Cao *et al*., 2019), while some scholars hold the opposite view (Li *et al*., 2018; Wan *et al*., 2018; Li, 2021b). I.R. Brooking (1990) investigated the relationship between dry matter and moisture in grain and ear of 18 maize cultivars from silking to harvest, and divided the process of grain dehydration into two stages. The water loss at the initial stage from grain filling to physiological maturity, was called as the developmental dehydration process related to grain filling. The second stage began with the physiological maturity to the water content of about 20%-25% suitable for grain harvesting, which was the physical dehydration process of grain drying. The research on the characteristics of grain dehydration by Cao *et al.* (2019) showed that the grain dehydration rate increased at first and then decreased, while at the later stage it increased rapidly. The grain dehydration rate displayed different characteristics before and after the starting point of rapid increase in the later stage, which was taken as the separation point. T1 stage was the early stage and T2 stage was the later stage. In T1 stage, grain dehydration rate was significantly positively correlated with grain filling rate *(P* < 0.01), but not significantly correlated with meteorological factors, indicating physiological dehydration properties. While in T2 stage, grain dehydration rate was strikingly positively correlated with temperature-related meteorological elements and average sunshine hours, and negatively correlated with rainfall, suggesting net dehydration characteristics. Thus far, people have only speculated on the mechanism of grain dehydration through grain development properties and correlation analysis of dehydration rate with grain filling rate and meteorological factors, which still lacks further evidence.

After pollination of the maize floret, the plant inputs nutrients and water into the developing grain through the pedicel vascular bundle. The pedicel exists at the base of the caryopsis and contains two outer phloem vascular bundles with a cluster distribution at the end (Zhang *et al*., 1999), which is branched from the longitudinal vascular bundles of the cob near the grain side. It is the direct supplier of grain nutrients (Shao *et al*., 2016). The water and nutrients in the pedicel vascular bundle can not be transported directly to the endosperm, but must first be unloaded by the placenta-chalazal layer (P-C) between the vascular bundle and the basal endosperm cells, before being transported to the endosperm (Kiesselbach, 1949; Balcon *et al*., 2004). Nutrients unloaded from vascular bundles to P-C region are transported through the endosperm nutrient transport tissue and stored in endosperm (Wang *et al*., 1997). In the process of grain filling, the migration pathway of water in grains is still controversial (Zhang *et al*., 2007). Some studies confirmed that the water in cereal caryopsis was mainly lost from the caryopsis pericarp by evaporation (Radley, 1976; Lee and Atkey, 1984). While other studies deemed that the water in caryopsis could be transferred to the maternal tissue (Meredith and Jenkins, 1975; Zhang *et al*., 2007).Previous studies suggested that physical damage or xylem blockage caused by growth pressure (growth strains) might result in reduction in xylem transport (Lang and Düring, 1991; Coombe and McCarthy, 2000; Dražeta *et al*., 2004; Knipfer *et al*.,2015). Also the excess water in the storage sink might be exported through the phloem, but separating the inflow of photosynthetic assimilates from the water outflow in the phloem became a problem (Patrick *et al*., 2001). Some researchers (e.g. Crafts and Currier, 1963) also put forward the “pressure flow hypothesis”, which held that a large number of assimilates were transported to the sink through the phloem with water as solvent, and if the water consumption (growth and transpiration) of the sink was not very large that the water transported through the phloem was able to meet or even exceed the demand, thus the surplus water was presumed to flow back to the parent tissue through the xylem. However, the pathway that water passes through the xylem in this process remains obscure (Zhang *et al*., 2007; Bewley *et al*., 2013a). Early studies had found that the water flowing into fruits or other storage organs through xylem was limited or even completely disappeared (Wiersum, 1966; Ollat *et al*., 2002) during the period of rapid growth and the fastest increase in volume. Xylem sap was the main water source of grape berries before veraison (discoloration, beginning of ripening, cell expansion after a short lag), while phloem sap became the main water source after veraison (Keller *et al*., 2006). At the onset of ripening, the decline of xylem inflow might be the result of an increase in sink-driven phloem influx (Keller *et al*., 2015). Zhang and Keller’s research revealed that a fraction of water imported berry through the phloem was used for berry growth and surface evaporation, and the surplus was recirculated via the xylem during grape berry development (Zhang and Keller, 2017), which was consistent with the “pressure flow hypothesis” proposed by Crafts and Currier (1963). A further research is still needed to determine whether the migration pathway of water in cob and grain accords with the viewpoint of “pressure flow hypothesis” in the process of maize grain filling and dehydration.

In order to demonstrate the previous inference about the driving force of grain dehydration, a grain coated technology was developed(patent number: 2018112795690). By means of the experimental operation of grain coated technology to prevent the water loss from the grain pericarp, the grain moisture content and dehydration rate of treatment and control during grain filling were compared to demonstrate whether grain filling and surface evaporation was the main dehydration driving force before and after grain physiological maturity, respectively. According to Keller’s series of studies (Keller *et al*., 2006; Keller *et al*., 2015; Zhang and Keller, 2017), this study referred to the method of soaking vines to transport apoplast dyes (basic fuchsin solution) into fruits, and continuously imported dyes from stems under ear-pedicel on living plants, then observed the movement of dyes in cob xylem. With reference to the method of transporting dyes into the fruit by soaking the grape fruit (the top was removed and exposing the xylem), the dye was continuously injected into the cob, and the movement of the dye in the cob, ear stalk and stem was observed. Meanwhile, the development dynamics of ear and grain were recorded, and the grain filling rate, dehydration rate, and contents of soluble sugar and starch in grain were determined. Combined with the results of dye movement experiment, the migration pathway of water in cob and grain were studied.

## Materials and methods

### Plant growth

The grain coated experiment was carried out in 2019 in Qingyuan Experimental Station of Hebei Agricultural University, Baoding City, Hebei Province, China (38°49’N, 115°26?; elevation 13m). The annual precipitation in the study year was 441.9mm, and the precipitation in the growing season was 344.8mm. The experimental materials were JNK728 (mid-maturing cultivar) and XY790 (early-maturing cultivar), which were sown on 15 June 2019. In the experiment, the planting area of each cultivar was about 200m^2^, the row spacing was 0.6m, and the designed planting density was 7.5×10^4^ plant hm^-2^. Controlled-release compound fertilizer (N-P_2_O_5_-K_2_O: 26-12-12) selected as base fertilizer was applied at sowing, and the rate of fertilizer application was about 750 kg hm^-2^. No-tillage before sowing summer maize, artificial sowing, 2 seeds per hole. Irrigating after sowing to ensure seedling emergence, then thinning seedling during the 4-leaf period. The management of irrigation, weeding and pest control during the growth period of maize were the same as those of the local production fields. The daily average temperature data of the test site were obtained from the small weather station (Watch Dog 2900 Weather Station) installed in the test site.

The dye movement experiment was conducted in the Innovation Experiment Park of Hebei Agricultural University, Baoding City, Hebei Province, China in 2021(38°49’N, 115°26?; elevation 13m). The experimental materials were two maize cultivars JNK728 (mid-maturing cultivar) and XY779 (early-maturing cultivar) and sweet maize cultivar WT2000. JNK728 sowed on 30 May, while XY779 and WT2000 sowed on 9 June. The planting area of each cultivar was about 100 m^2^, and the cultivating practices were same as the grain coated experiment excepting irrigation. After irrigating after sowing, irrigation was executed by controlling the lower limit of soil moisture content. Irrigation begins when the soil moisture content is reduced to 70%, and the upper limit is 100%. The temperature data in the dry shed were recorded by the temperature recorder (Microlite USB DATA LOGGERS, LITE5032L).

### The grain coated experiment

The plants with the same growth were marked in each plot of the two cultivars in the silking stage in the field. Starting from 35 days after pollination (DAP), ear husk of three marked plants selected from each cultivar plot were carefully stripped down layer by layer until the grain of middle and upper ear position was exposed, and the husks were not hurt as far as possible. The central portion of the ear was used as the experimental treatment area. Tweezers were used to remove the grains of experimental treatment area in alternate rows and alternate grains, which was convenient for the coated treatment of the remaining grains. Maintain the intactness of the grains used for coated treatment during operation, so that they can still grow normally. After completing the grain removal operation, the grain was coated with a quick-drying liquid material which is no effect on the grain development. The coating dries quickly, then forms a waterproof and tough film that tightly wraps the grain and prevents the water in grain from evaporating through the pericarp. In the experimental treatment area of ear, half of the remand grains were coated, and the other half uncoated were control. When the treatment was completed, the husks were restored layer by layer from the inside to the outside, and the whole treated ear was bound with string to maintain the original posture of the husks. In order to avoid the grain mildewing caused by the rain entering from ear top, the husk of ear top was capped with adhesive tape (Fig. 1). The ears were treated per 5 days, and the last treated ears were brought back to the laboratory for further analysis until they were harvested after physiological maturity. Tearing off the waterproof coating outside the treated grains on the ear, 20 grains were gained from treatment and control respectively of each ear to determined the fresh weight (FW) and volume, and then were dried in a constant temperature oven at 85 °C to constant weight and were recorded as the dry weight (DW). Grain filling rate(GFR), water content(WC), moisture content(MC) and dehydration rate(DR) were calculated according to grain fresh weight and dry weight as follows:

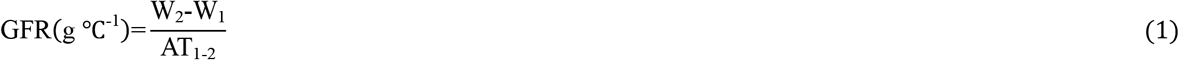

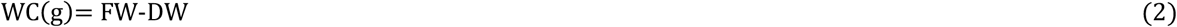

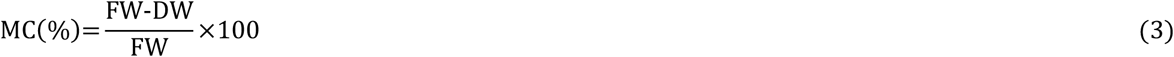

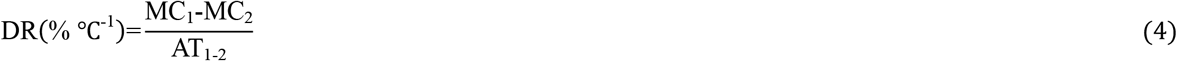

where W_1_ and W_2_ is the dry weight of previous sample and the last sample, respectively; AT1-2 is the accumulated temperature between two samples; MC_1_ and MC_2_ is the moisture content of previous sample and the last sample, respectively.

**Fig. 1.**
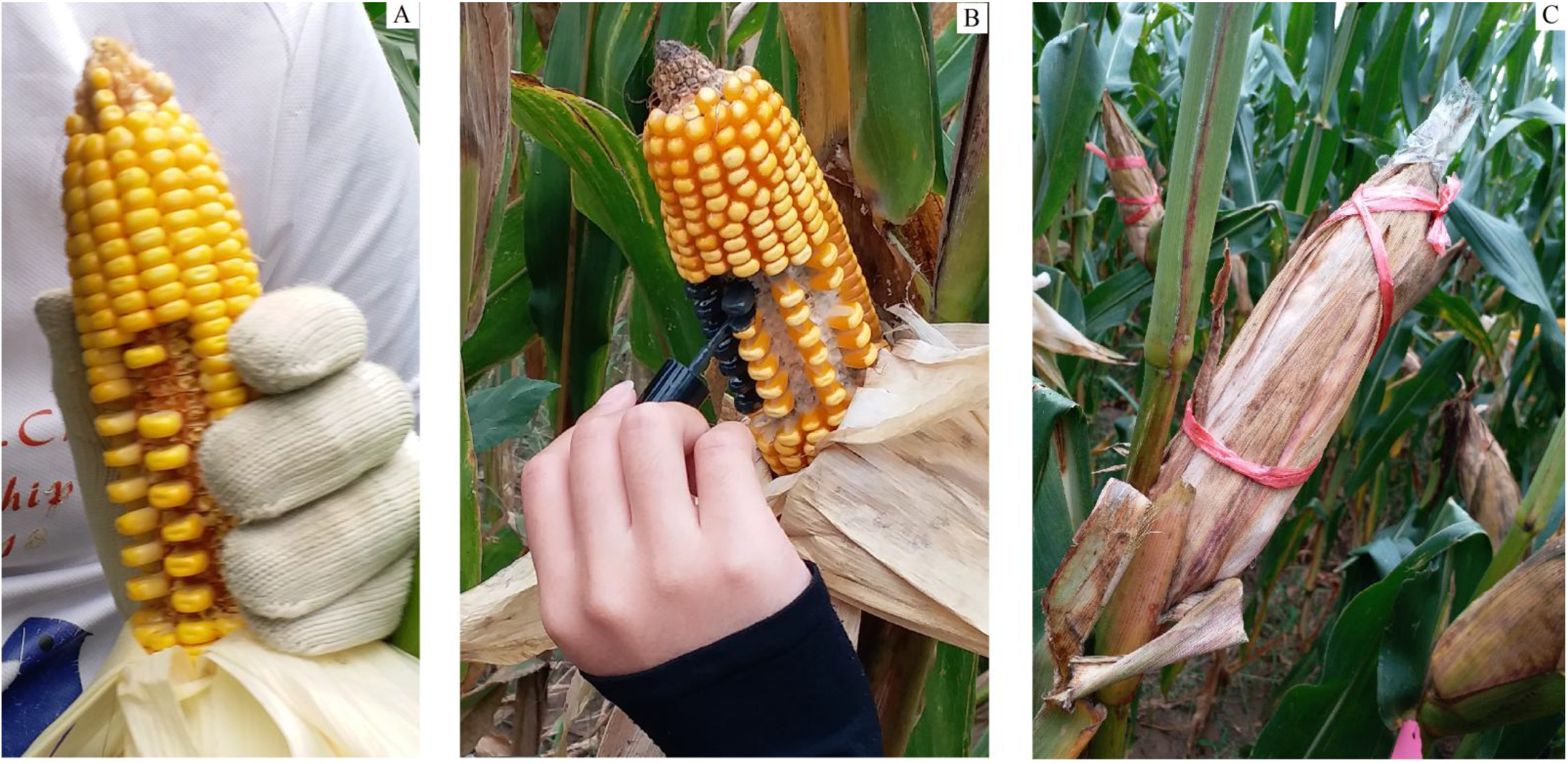
The operational processes of grain coated experiment. After taking off husk, grains at the middle of the ear were removed in alternate rows and grains(A), then coated the half of remaining grains(B), finally closed husk and bound the ear(C).

### The dye movement experiment

The plants with the same growth were marked in each plot of JNK728, XY779 and WT2000 during the silking period in the dry shed of the Innovation Experiment Park. Starting from 5DAP, 6 labeled plants were selected from each cultivar planting plot (The each treatment of dye input from stem and cob required 2 plants, and the remaining 2 plants were control). The 0.1% xylem-mobile dye (basic fuchsin solution, C20H19N3HCltermol.wt338Dqa) in an infusion bag, was continuously injected from stem attached to the ear-pedicel from below and from the cob (Fig. 2), and the infusion speed was adjusted to about 1drop per 15 seconds. After the infusion lasted for 2 days, the ear of control and dye infused from stem, the whole plant of dye infused from cob were brought back to the laboratory for analysis. It was treated per 5 days until harvested after physiological maturity.

**Fig. 2.**
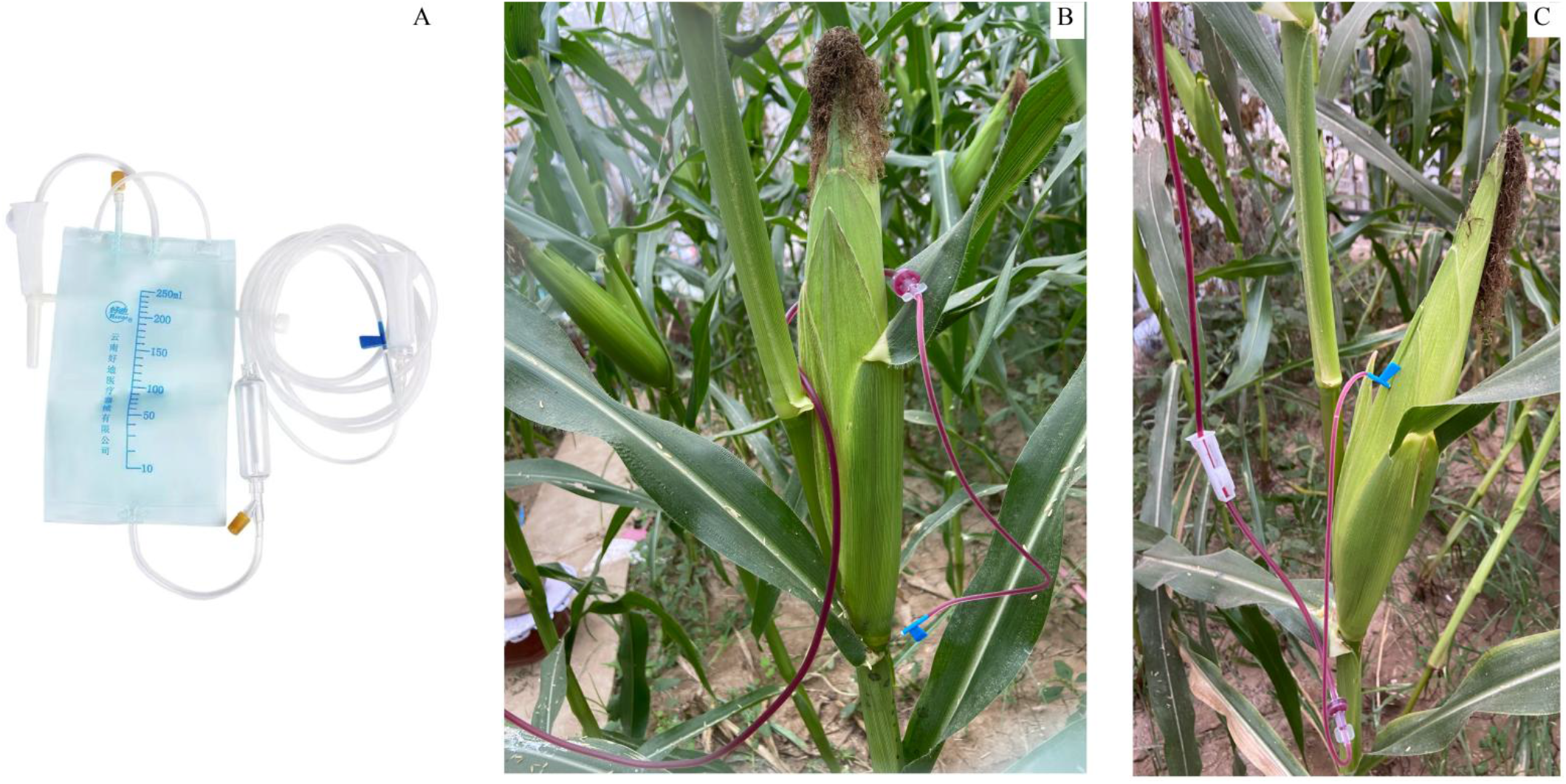
Dye movement experiments that the xylem-mobile dye basic fuchsin infused from the stem attached to the ear-pedicel from below(B) and from the ear cob(C). A is the infusion bag for the dye.

In the laboratory, the ears infused dye from stem were dissected longitudinally to observe the dye movement in the cob xylem. The ear, ear-pedicel and stem of the dye movement experiment that dye was infused from cob, were dissected longitudinally to observe the dye movement in the xylem of cob, ear-pedicel and stem. After breaking down the ears of two control plants to organs, the fresh weights of husks, 100 grains in the middle of the ear and cob (the cob started from 15DAP) were weighed, and the 100 grain volume was measured. Each organ was dried to constant weight at 85 °C in a constant temperature oven, and dry weight was recorded. The grain water content, moisture content, dehydration rate and grain filling rate were calculated according to the dry and fresh weight of each organ, and the calculation method is same to the grain coated experiment. The dried seeds were crushed by RS-FS1406, and then passed through a 100-mesh sieve. According to the national standard GB-5006-85, the contents of soluble sugar and starch in grains were determined by anthrone colorimetry (Wang, 2021). WT2000 only measured the grain dry weight, soluble sugar and starch content at 20 and 30DAP.

### Statistical analysis

Statistical analyses were carried out using Predictive Analytics Software (PASW) version 26.0 (IBM SPSS Statistics). The grain dry weight, volume, moisture content, filling rate and dehydration rate between treatment and control in grain coated experiment were analyzed in Paired-Samples T Test. In dye movement experiment, Pearson correlations were calculated to identify interrelationships among grain filling rate and grain dehydration rate, grain moisture content and husk moisture content, husk looseness, and grain dehydration rate and husk dehydration rate, husk loosing rate, respectively.

## Results

### The grain coated experiment

From 35 DAP to physiological maturity (Fig. 3), the JNK728 and XY790 grain dry weight and volume of coated treatment were smaller than that of control, but the difference was not significant, indicating that coated treatment had no effect on grain development. After physiological maturity, the grain dry weight of the two cultivars was similar to that of control, and the grain volume of treatment was larger than that of control, but the difference was not significant. Before physiological maturity, there was no difference in grain moisture content of middle maturing cultivar JNK728 between treatment and control, but the difference reached significant or very significant level after physiological maturity (Fig. 3A). For early maturing cultivar XY790, the grain moisture content of treatment was significantly or very significantly higher than that of control from 5 days before physiological maturity (Fig. 3B).

**Fig. 3.**
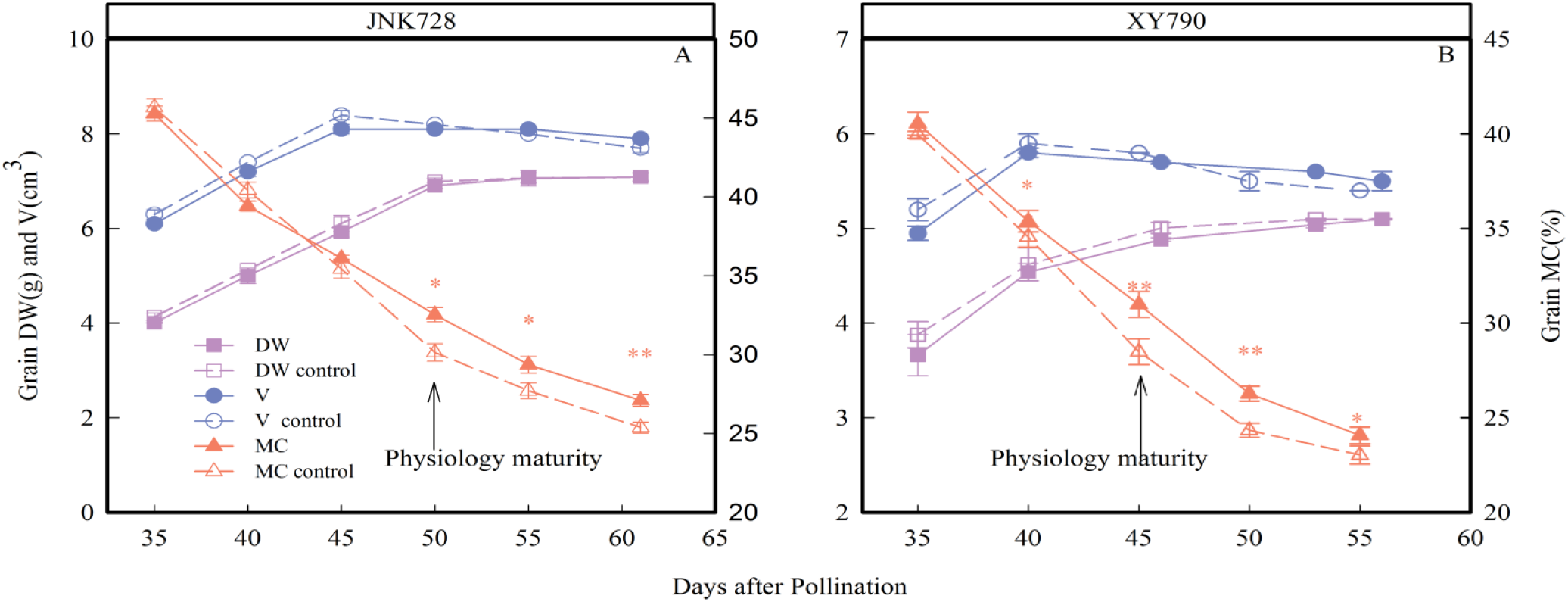
Dry weight (DW), volume (V) and moisture content (MC) of JNK728(A) and XY790(B) 20 grains between the late stages of grain filling. Paired-Samples T Test was performed for treatment and control data, * indicates significance at the 0.05 probability level, ** indicates significance at the 0.01 probability level. Values are means ± SE (*n*=2).

The grain filling rate of JNK728 and XY790 treatments was lower than that of control, but the difference was not significant. The grain filling rate of JNK728 was higher than that of XY790 (Fig. 4). The grain dehydration rate of JNK728 and XY790 was significantly or very significantly lower than that of control from 5 days before physiological maturity (45DAP) and 10 days before physiological maturity (35DAP) to harvest (Fig. 4), respectively. This experiment demonstrates the previous corollary (Brooking, 1990; Cao *et al*., 2019) that grain dehydration was divided into two stages, which was mainly driven by grain filling and surface evaporation, respectively. The grain filling rate, dehydration rate and moisture content of early and middle mature maize cultivars were not significantly different from those of control before 5-10 days before physiological maturity, and this dehydration process is driven dominantly by grain filling. When the grain development is close to physiological maturity, the grain filling rate decreases to the lowest, and there are striking or very striking differences in dehydration rate and moisture content between treatment and control. In addition, in this phase grain dehydration is driven dominantly by surface evaporation. When grain approaches physiological maturity, the grain of early-maturing cultivar loses water via surface evaporation as the main dehydration pathway earlier than the middle-maturing cultivar.

**Fig. 4.**
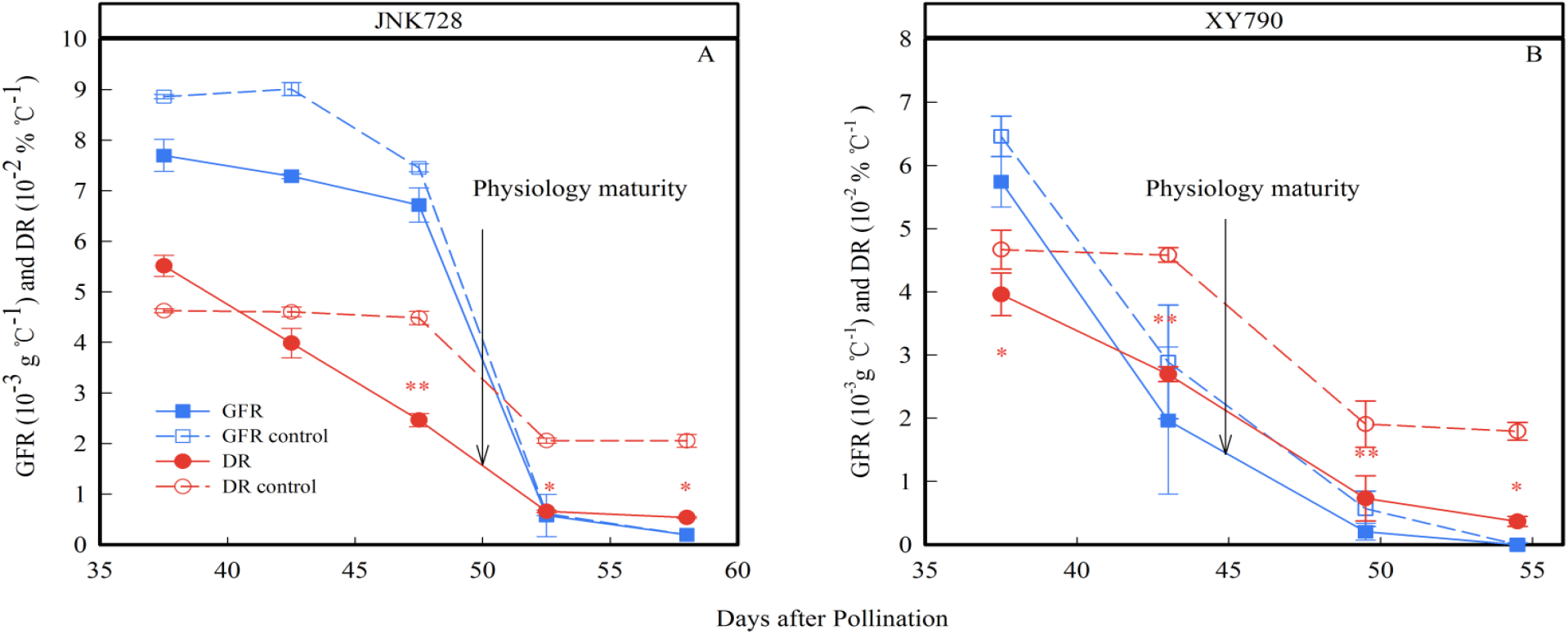
Grain filling rates (GFR) and dehydration rates (DR) of JNK728 (A) and XY790 (B) 20 grains at the late stages of grain filling. Paired-Samples T Test was performed for treatment and control data, * indicates significance at the 0.05 probability level, ** indicates significance at the 0.01 probability level. Values are means ± SE (*n*=2).

### The dye movement experiment

Dye was infused from stem attached to the ear-pedicel from below. For JNK728 and XY779, the dye could move through the pedicel xylem below the grain within 10 days after pollination (Fig. 1A, B). The dye was not observed in pedicel at 15 DAP (Fig. 5C), but in the longitudinal vascular bundles of cob. In the stage of grain formation, the grain absorbed water rapidly and expands its volume (Borrás and Gambin, 2010). In the process that plant transports nutrients and water to the ear, part of the water as the solvents of nutrients enters the grain through the pedicel phloem, and other part of the water enters the grain through the pedicel xylem (Westgate and Boyer, 1984), thus basic fuchsin dye could be observed in the cob pedicel at this stage. These indicate that the cob transports water to the grain no longer through the pedicel xylem, but through the phloem from 15 DAP. That is to say, phloem is the unique path that water enters grain after 15 DAP. Almost in 25-30 DAP, the dye only remained confined to the ear stalk or the bottom of cob (less entered the cob), then the dye could move throughout the xylem of cob again from 35 DAP (Fig. 5D). The results show that water can not enter the ear via the cob xylem, and the growth and development of the ear is mainly maintained by the water import via the cob phloem during 25-30 DAP. The dye infused from the stem could also move throughout the pedicel xylem of sweet maize WT2000 at 20 DAP (Fig. 5E). Furthermore, the dye in the pedicel xylem of the cob bottom and apex position was darker than that of the cob middle position, then the dye was not observed in pedicel from 25 DAP. The pedicel dyed time of WT2000 was longer than that of JNK728 and XY779 (later than that of JNK728 and XY779). In other words, water was continuously transported to grain of WT2000 through the phloem and xylem of pedicel from pollination to 20 DAP. Due to the dye in the pedicel xylem of the cob middle position was lighter than that of the cob bottom and apex positions, it is speculated that the water transported through the pedicel xylem to the grain of the ear middle position is less than that of the ear bottom and apex position.

**Fig. 5.**
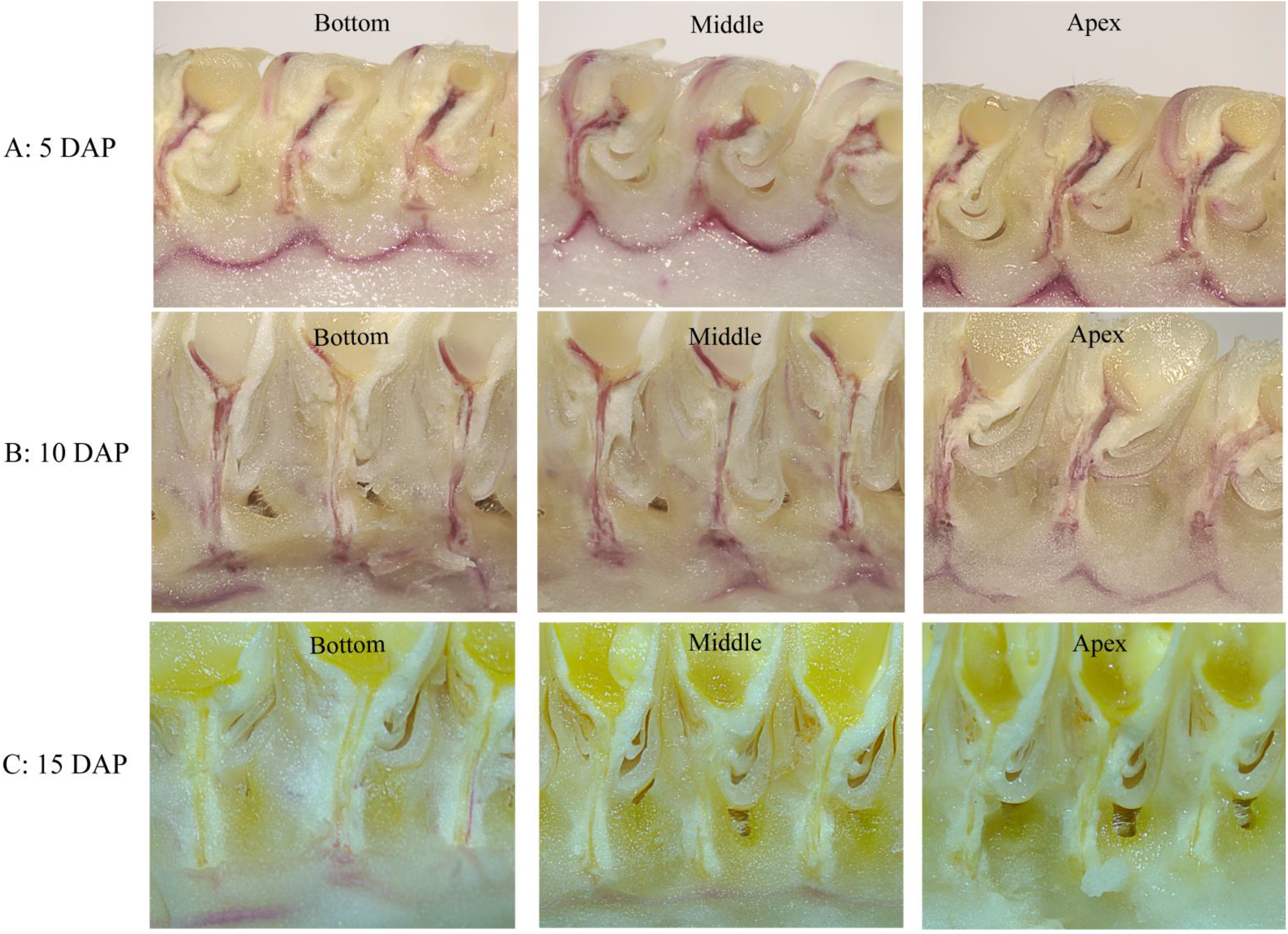
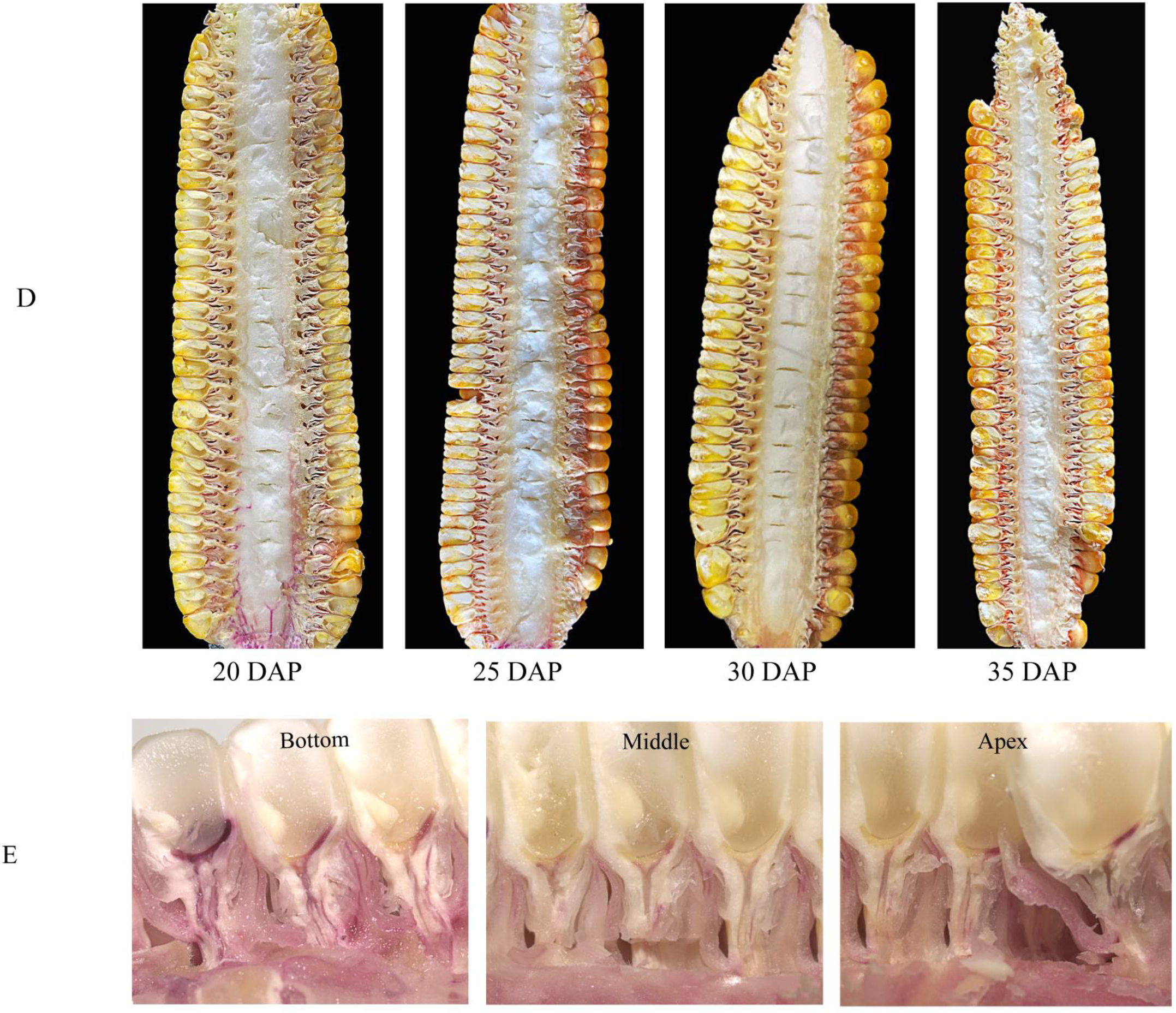
Movement of the xylem-mobile dye basic fuchsin infused through the stem attached to the ear-pedicel from below. Cross-section of the ear different position at 5 DAP (A), 10 DAP (B) and 15 DAP (C) for JNK728, and 20 DAP for WT2000 (E). Cross-section of the ear from 20 DAP to 35 DAP for JNK728 (D).

Dye was infused from cob. The dye remained almost restricted in the cob, and no traces of dye were observed in the ear-pedicel and stem at 5 DAP(during the period of grain formation) (Fig. 6A, B), then the dye was observed in cob, ear-pedicel and 2 stems attached to the ear-pedicel from 15 DAP(Fig. 6C, D). It is speculated that there is only the net import of water via cob xylem in the ear during the grain formation period. Combined with the experimental results that dye was infused from stem, it is inferred that there is only net import of water via cob xylem in the ear from pollination to grain formation, both import and export from grain formation to 25 DAP and from 30 DAP to physiological maturity, and net export in 25-30 DAP. The surplus water flowed back to the plant from ear through the xylem and was recycled by the plant during the development of ear. From being able to transport water into the ear smoothly, the xylem of cob experiences the processes of transport stop and resuming, along with transporting the surplus water from ear back to the plant. The results prove that the xylem of cob ceases importing water to the ear not because the xylem vessel is blocked, but because the direction of water transport in the xylem of cob changes from towards ear to towards plant. By the same token, it can also be speculated that the reason why the cob no longer transports water to the grain through the pedicel xylem is that the surplus water in grain is transported back to the cob via the pedicel xylem, instead of the blockage of the xylem vessel.

**Fig. 6.**
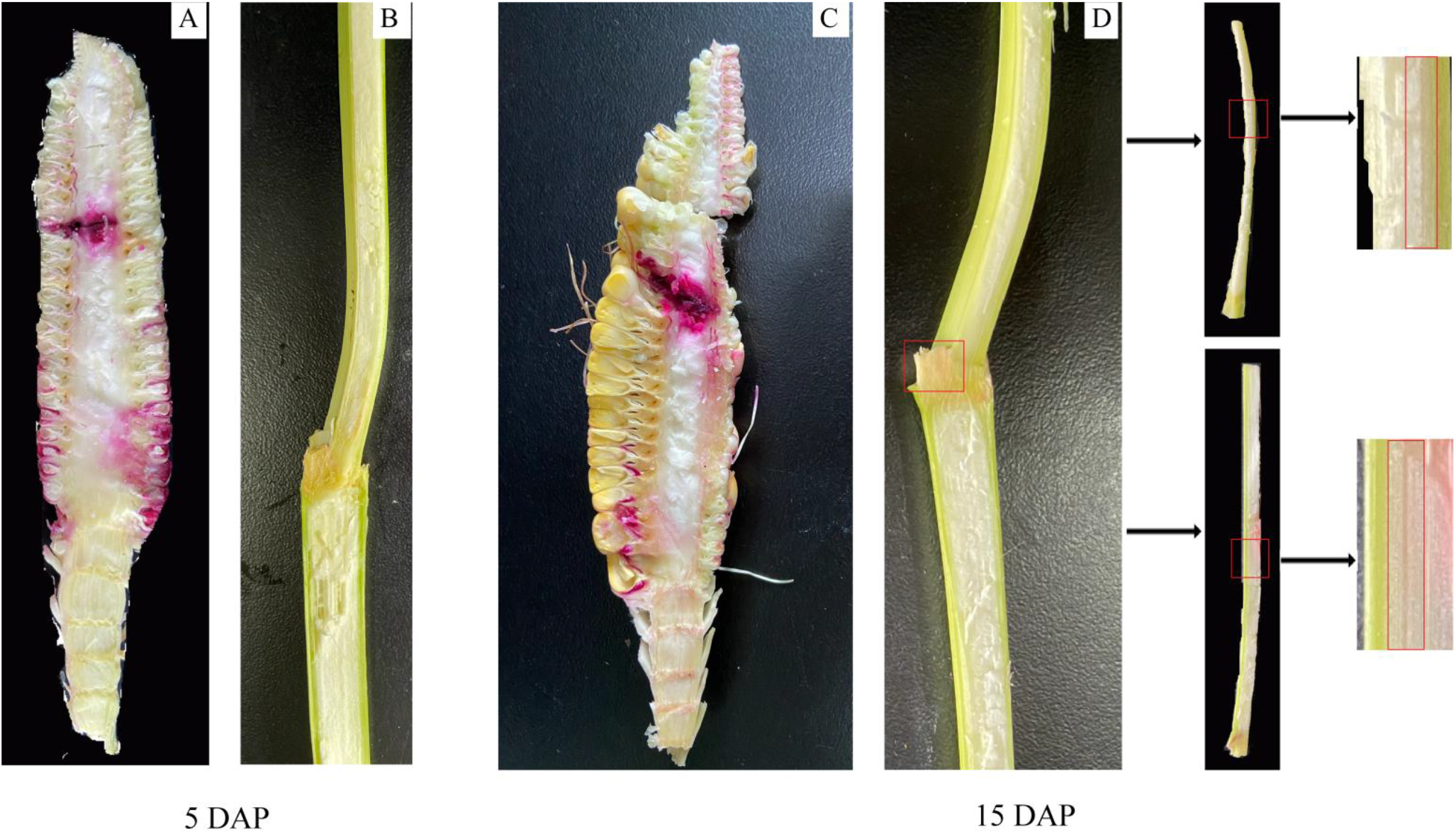
Movement of the xylem-mobile dye basic fuchsin infused through the cob of JNK728 ear. Cross-section of the ear at 5 DAP (A) and 15 DAP (C), and of the 2 stems attached to ear at 5 DAP (B) and 15 DAP (D).

Fig. 7 showed the daily average temperature in the experimental garden after maize pollination. The water content of cob and 100 grains of JNK728 and XY779 reached the maximum at 20 DAP (Fig. 8A, B), and the volume of 100 grains reached the maximum at 35 DAP (Fig. 8C), so the ear needed less water during subsequent development.

**Fig. 7.**
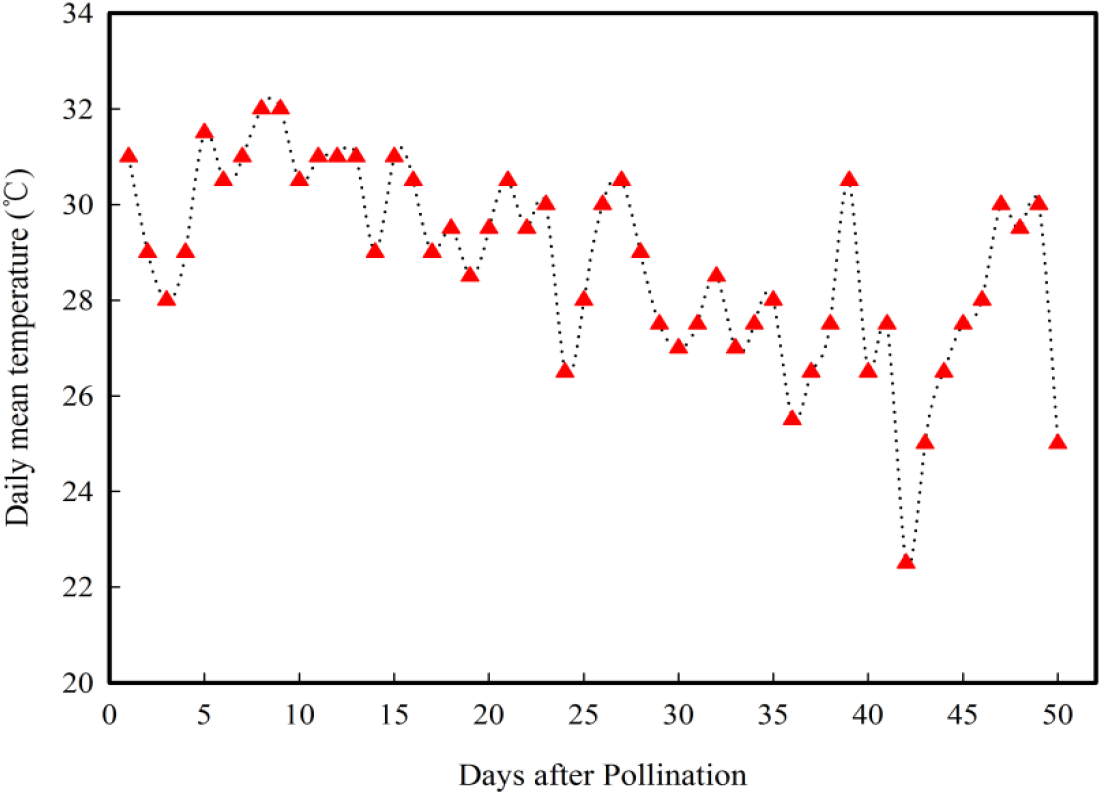
Daily mean temperature data during grain filling stage.

**Fig. 8.**
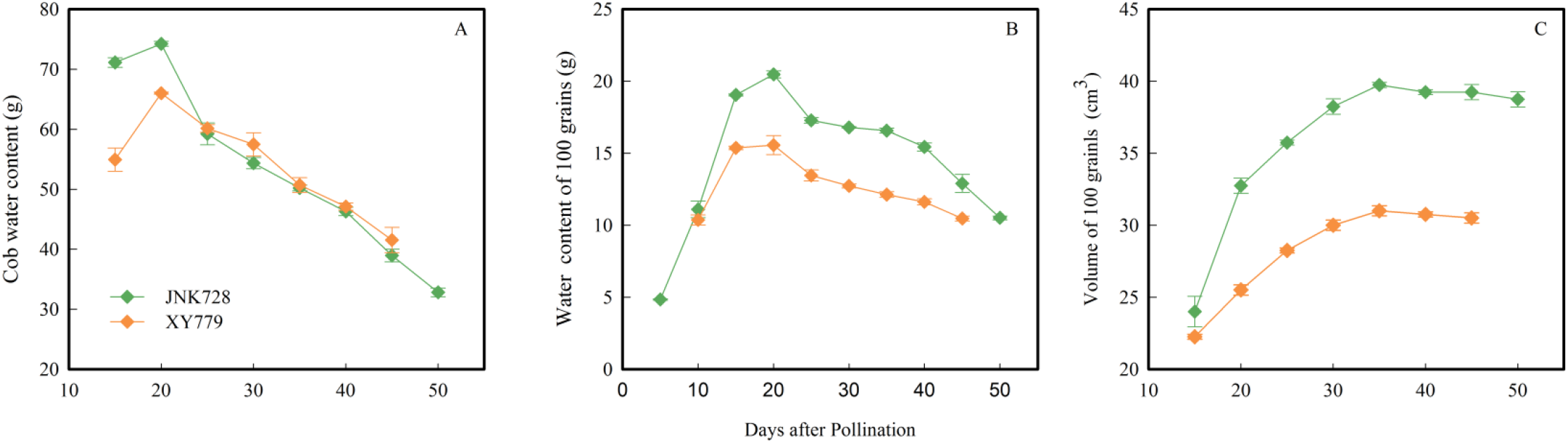
Water content of 100 grains (A) and cob (B), and 100 grains volume (C) of JNK728 and XY779 during grain filling stage. Values are means ± SE (*n*=2).

The soluble sugar content of two maize cultivars of JNK728 and XY779 showed a bimodal trend (Fig. 9A, B), and consistently reached the maximum at 10 DAP, then decreased. The soluble sugar content of JNK728 slowly increased from 25DAP to the second peak in 35 DAP, and that of XY779 increased to the second peak in 30 DAP. The soluble sugar in the grain accumulated rapidly from pollination to 10 DAP, meanwhile the osmotic pressure increased and the water potential decreased in grain(Westgate and Boyer, 1986), which promoted more water to enter the grain through the pedicel phloem and xylem. The starch content of the grain began to increase rapidly in 10-15 DAP, and simultaneously, the soluble sugar content of the grain decreased sharply, suggesting that the sugar began to convert rapidly into starch and the filling rate accelerated. After the formation of grain, the grain filling rate began to accelerate, so did the phloem transport rate (Patrick and Offler, 1995; Walker *et al*., 2000). The water entering the grain via the pedicel phloem could gradually meet or even exceed the needs of grain development. The pedicel xylem gradually reduced and finally stopped transporting water to the grain, and even transported the surplus water in grain back to the cob. The soluble sugar content of sweet maize WT2000 in grain decreased gradually from the bottom to the apex of the ear, and the starch content in grain of the ear middle position was higher than that of the ear bottom and apex position at 20 DAP (Fig. 9C). In other words, the grain filling rate of the ear middle position was faster than that of the ear bottom and apex position. The amount of water entering the grain of the ear middle position via the pedicel phloem increased, as a result that the water importing the grain of the ear middle position through the pedicel xylem decreased, which was consistent with the results of the dye movement experiment (Fig. 5E).

**Fig. 9.**
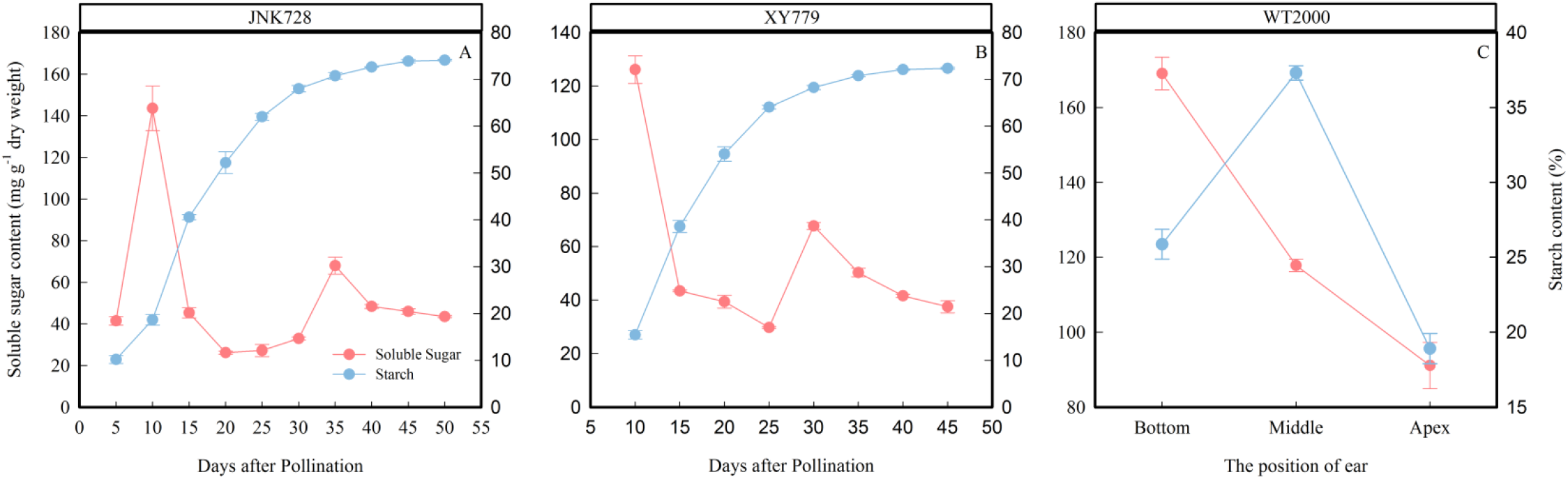
Soluble sugar content and starch content of JNK728 and XY779 grains during grain filling stage, and ET2000 grains of ear different position at 20 DAP. Values are means ± SE (*n*=2).

The curve chart of grain filling rate and dehydration rate of two maize cultivars of JNK728 and XY779 showed that the change trend of grain filling rate and dehydration rate was very close, with the maximum value in 15-25 DAP (Fig. 10). A striking positive correlation was observed between grain filling rate and dehydration rate from grain formation (after 10 DAP) to physiological maturity (Fig. 11). Starting from 30 DAP, the husk withering and drying process, moisture content and loose degree of XY779 was faster, lower and higher than that of JNK728 (Fig. 12, Fig. 13), respectively. The husk of JNK728 and XY779 loosened rapidly in 35-40 DAP and 30-35 DAP, respectively. At the same time, there was a very striking positive correlation between husk and grain moisture content (Fig. 14A), and a very significant inverse correlation between husk loose degree and grain moisture content (Fig. 14C). The correlation between husk and grain dehydration rate of JNK728 was inversely significant, but that of XY779 was not marked (Fig. 14B). There was no correlation between husk loosening rate and grain dehydration rate (Fig. 14D). From grain formation to 35 and 30DAP for JNK728 and XY779 respectively, the grain filling rate of was relatively high. The husk was not completely withered and dried, and moisture content was high, so that the ear was wrapped tightly by husk at this phase. The driving force of grain dehydration at this stage was grain filling, which was also demonstrated by the correlation analysis between grain filling and dehydration rate. After the rapid decrease of grain filling and dehydration rate, the moisture content of husk decreased rapidly to less than 40% at 40 DAP, and the husk of JNK728 and XY779 withered completely and loosened rapidly at 40 and 35 DAP, respectively. Although the grain filling rate continued to decrease at this stage, the dehydration rate remained stable, and still slowly increased with the progress of development. The results show that the grain filling rate decreases rapidly as grain filling approaches the end, the effect of grain filling driving dehydration weakened. Along with the withering and loosening and the decrease of moisture content of husk, the main driving force of grain dehydration gradually changes from grain filling to surface evaporation. Combined with the results of grain coated experiment, it is proved that from grain formation to approximated physiological maturity (5-10 days before physiological maturity), the main driving force of grain dehydration is grain filling, and then changes to surface evaporation.

**Fig. 10.**
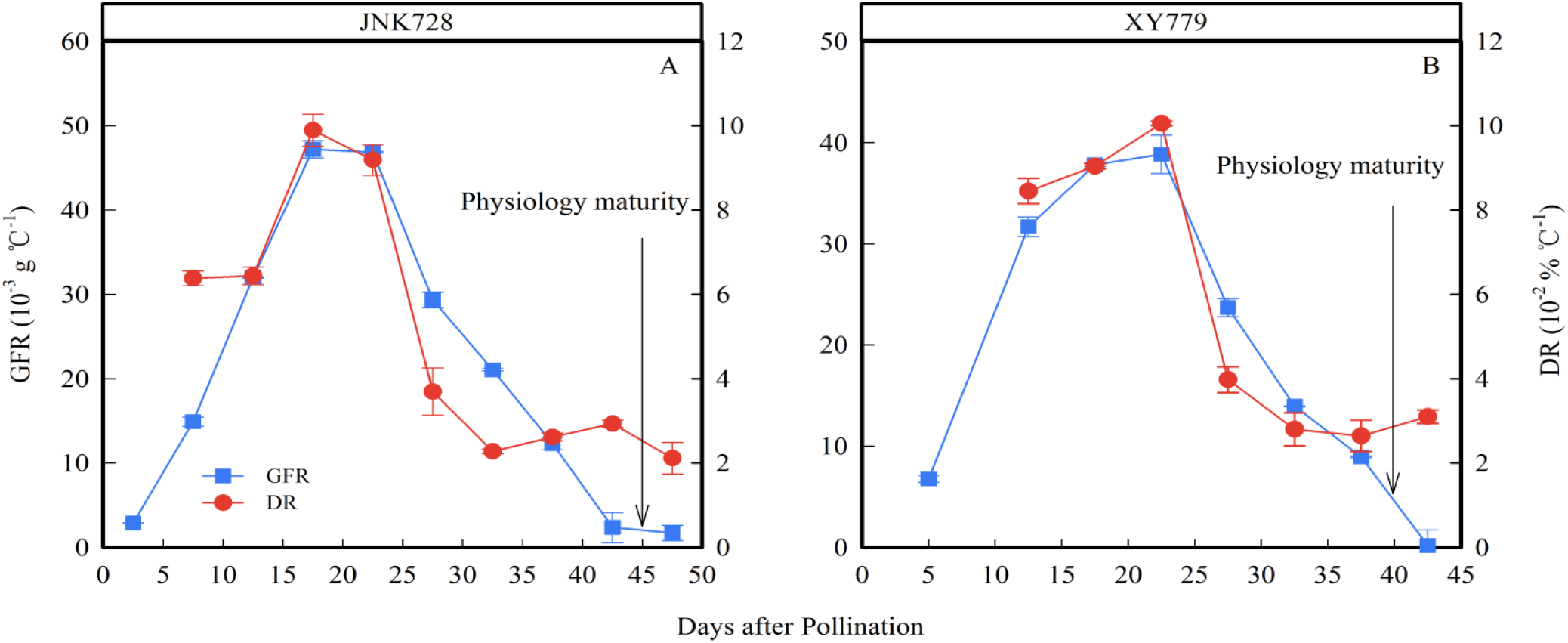
Grain filling rates (GFR) and dehydration rates (DR) of JNK728 and XY779 100 grains during grain filling stage. Values are means ± SE (*n*=2).

**Fig. 11.**
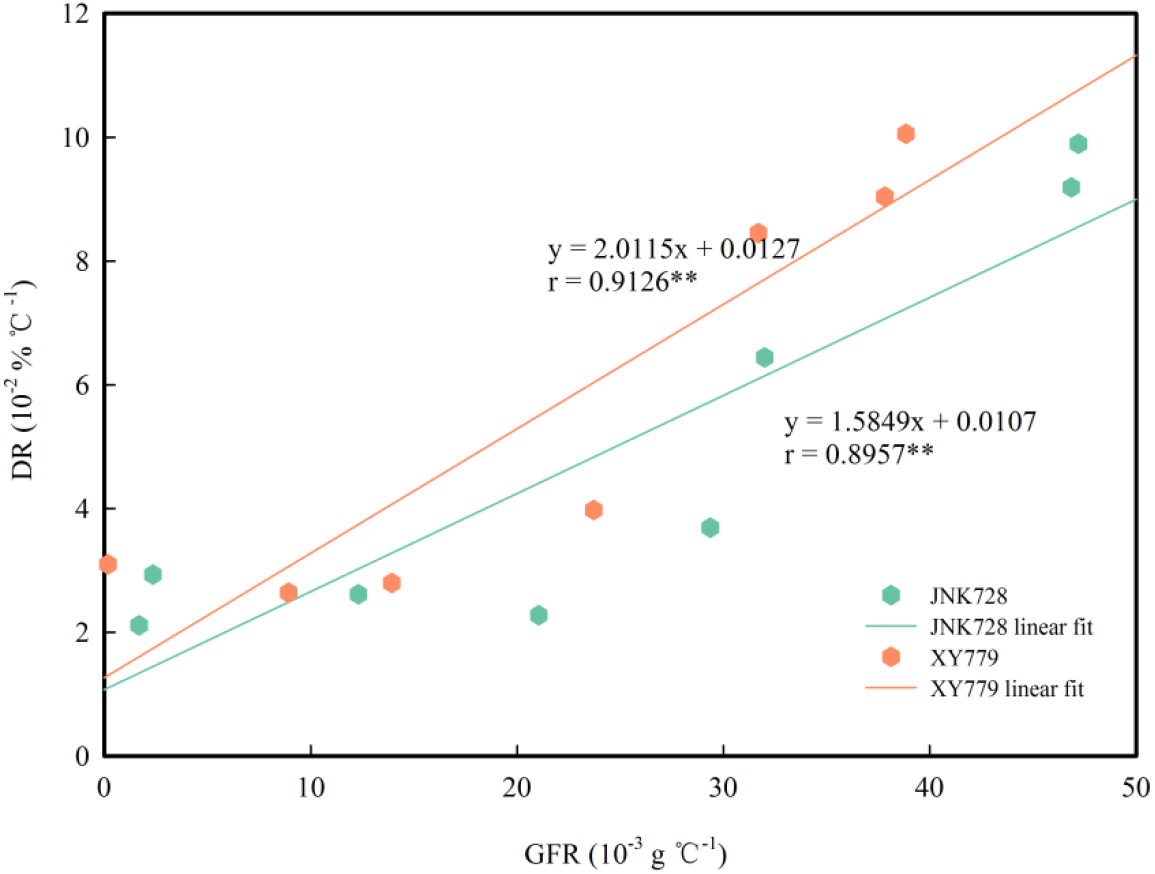
Relationship between grain filling rates (GFR) and dehydration rates (DR) of JNK728 and XY779 during grain filling stage. ** indicates significance at the 0.01 probability level.

**Fig. 12.**
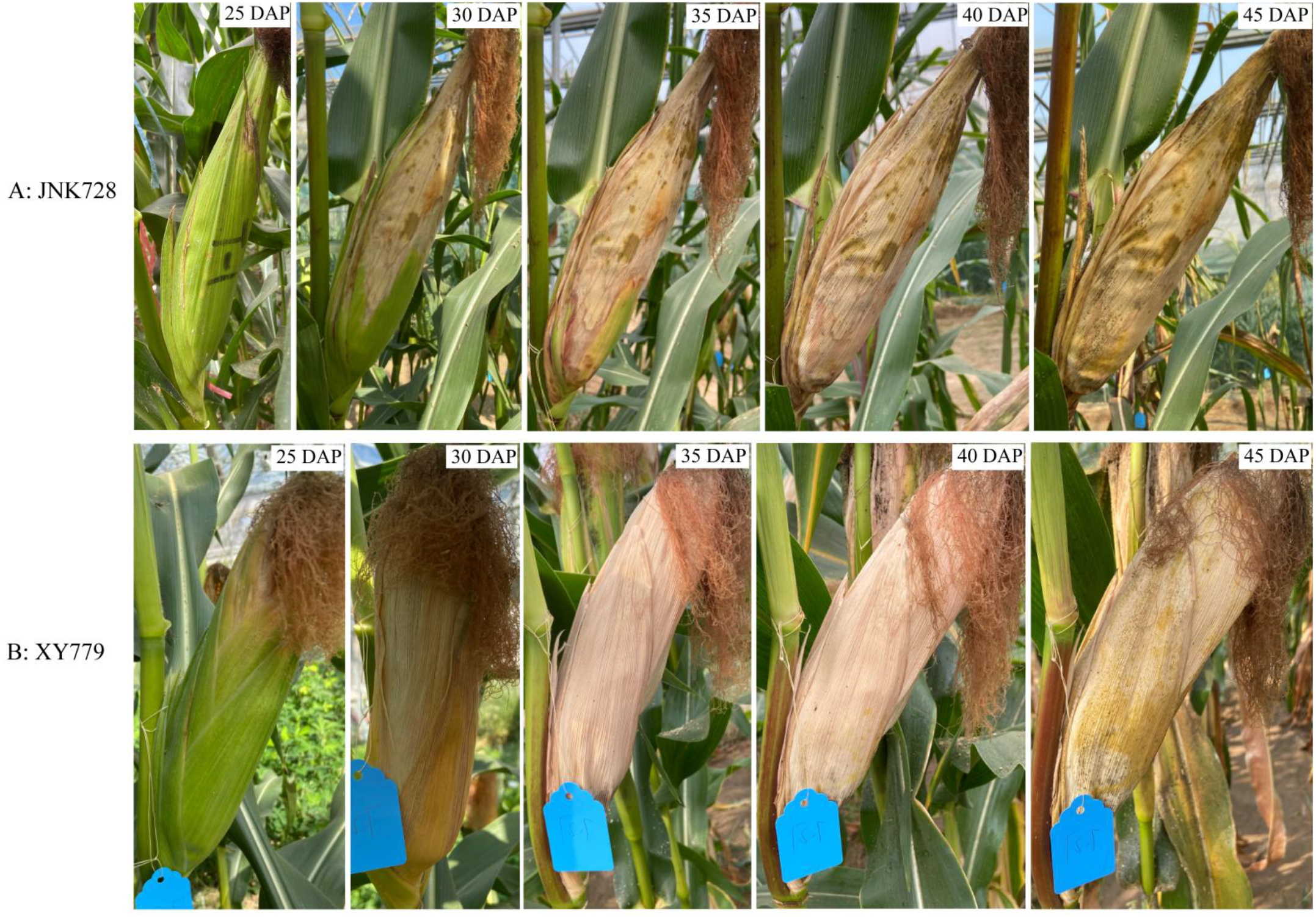
The husk withering progress of JNK728 and XY779 from 25 to 45 days after pollination (DAP).

**Fig. 13.**
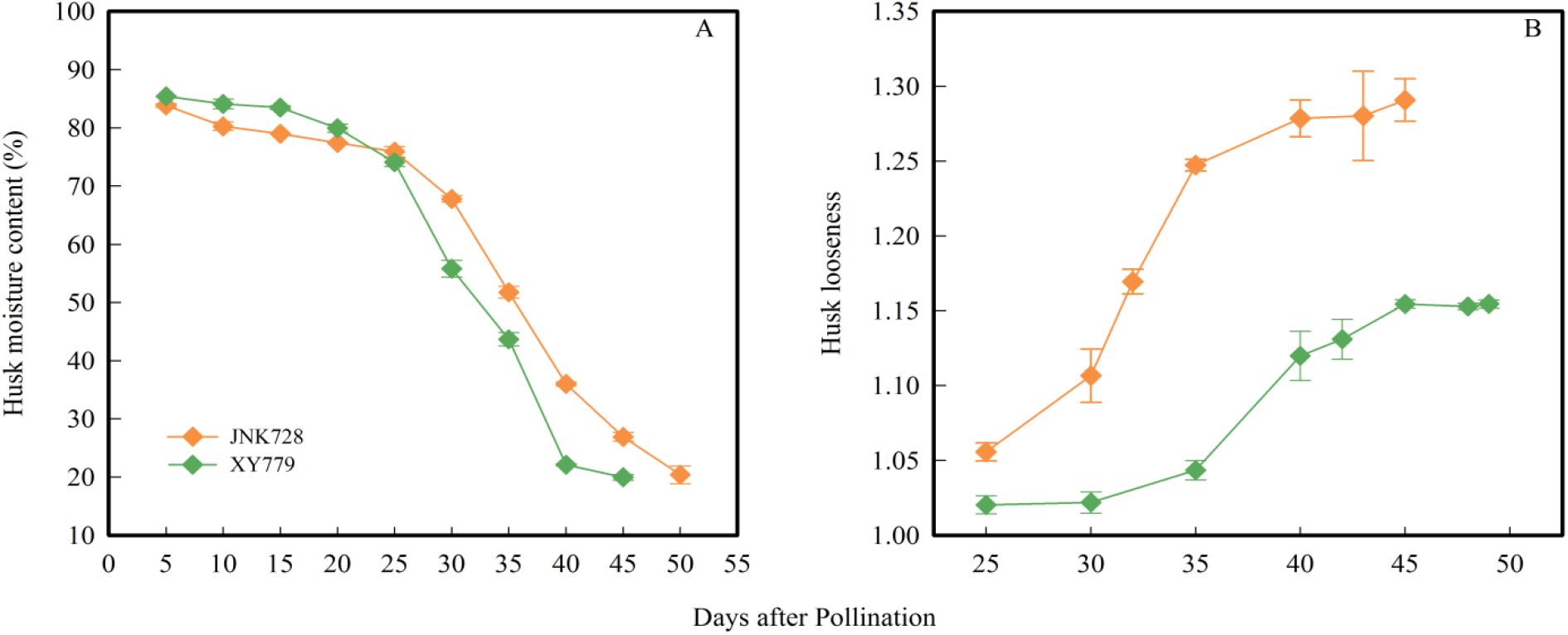
Husk moisture content and looseness of JNK728 and XY779 during grain filling stage. Values are means ± SE (*n*=2).

**Fig. 14.**
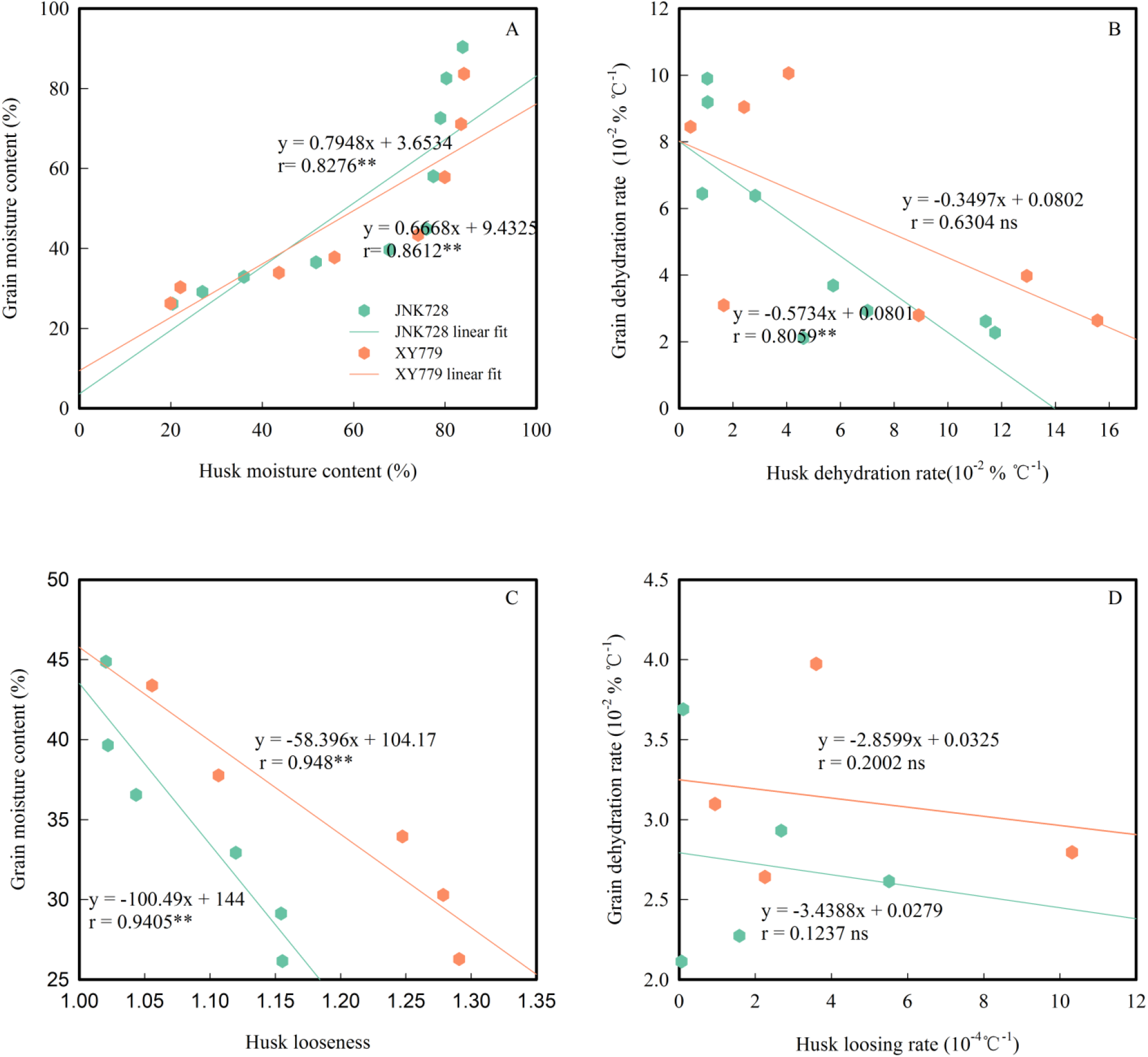
Correlation analysis between grain and husk moisture content (A), between grain and husk dehydrating rate (B), between grain moisture content and husk looseness (C), and between grain dehydrating rate and husk loosing rate (D) of JNK728 and XY779 during grain filling stage. ** indicates significance at the 0.01 probability level, “ns” indicates no significant.

## Discussion

The grain removing before coated treatment magnified the grain growth space in the experimental treatment area on the ear. In the process of grain filling, the coated grains were wrapped by the waterproof membrane, which had flexibility, but limited the increase of grain volume to a certain extent. By contrast, the change of control grain volume was not limited by space. Shen *et al.* (2019) pointed out that the grain weight of incomplete pollination treatment was larger than that of synchronous pollination treatment, indicating that after reducing the number of grains, the remaining grains had more nutrients and space for growth. Our measurements of grain dry weight, volume and filling rate support this conclusion. Before physiological maturity, the grain dry weight, volume and filling rate of JNK728 and XY790 of coated treatment were lower than that of control, but the difference was not significant, indicating that coated treatment had no effect on normal grain growth (Fig. 3, Fig. 4).

From roughly 5 days (45 DAP) and 10 days (35 DAP) before physiological maturity to after physiological maturity for JNK728 and XY790 respectively, the grain dehydration rate of the 2 cultivars was significantly or very significantly lower than that of control (Fig. 4), and the main driving force of grain dehydration changed from grain filling to surface evaporation. From 35 DAP to physiological maturity, the grain filling rate of XY790 was lower than that of JNK728. Furthermore, the grain filling rate of XY790 at 10 days before physiological maturity was lower than that of JNK728 at 5 days before physiological maturity (Fig. 4). In the later stage of grain development, the dominance of grain dehydration of XY790 transferred from grain filling to surface evaporation earlier than that of JNK728. Exactly as showed in Fig. 4, the occurrence time of significant difference in grain dehydration rate between treatment and control of XY790 was five days earlier than that of JNK728. The results of this experiment are a little different from Brooking’s and Bewley’s research (Brooking, 1990; Bewley *et al*., 2013b). They took the physiological maturity as the demarcation point of different dehydration stages, but this demarcation point was advanced in present study. The demarcation point of middle-maturing cultivar JNK728 and early-maturing cultivar XY790 was advanced about 5 and 10 days, respectively. Because Cao *et al*. (2019) studied the spring maize cultivar, the light and thermal resources were still sufficient to accelerate the dehydration rate in the late filling stage. However, summer maize cultivars were studied in this research, and there was little room for the increase of grain dehydration rate due to the lack of accumulated temperature in the late filling stage, so the results of this experiment were also different from those of Cao.

Basic fuchsin solution was continuously injected into JNK728, XY779 and WT2000 from the stem attached to the ear-pedicel from below. Seed development formed embryo and endosperm tissue through cell division and tissue differentiation, then water inflow drived cell expansion, and then dry matter began to accumulate (Bewley *et al*., 2013a). Nutrients (assimilates) and water transported from the maternal plant to the P-C (placenta-chalazal) region through the pedicel vascular bundle, are released to the extracellular space (apoplast) between the maternal and filial tissues, and then are imported to the developing endosperm and embryo. In the early stage of seed development, the phloem conductivity providing nutrients for seed is low (Zhang *et al*, 2007). The water entering the grain through the phloem can not meet the needs of grain development, and the xylem also provides water for the early expansion and growth of the grain (Westgate and Boyer, 1984). Therefore, in this study, the dye could enter the P-C zone following water through the pedicel xylem during the grain formation period (Fig. 5 A, B and E). High concentration photoassimilates (sucrose and other assimilates) are loaded into the source phloem cells that produce osmotic potential gradients, thereby attracting water to the cells, and increasing the turgor pressure of the phloem at the source. Which results in a driving force, namely hydrostatic pressure gradient, to transport the photoassimilate solution to the sink organs (van Bel, 1996; Lalonde *et al*., 2003; Koch, 2004; Turjeon, 2010; Patrick, 2013). In sink tissue, sucrose unloading from the phloem decreases the solute concentration in the sieve tube, resulting in water diffusing out of the sieve tube into surrounding cells and lowering the pressure, and further propelling the bulk flow of solutes (sucrose and other assimilates) from source to sink tissues (Lalonde *et al*., 2003; Patrick, 2013). At the stage of grain formation (within 10 DAP), the content of soluble sugar in grains increased rapidly (Fig. 9A, B). The grain filling rate is low, as results that the starch accumulation is less and the solute in grain is mainly soluble sugar, causing elevating the osmotic pressure and lowering the water potential in grain (Westgate and Boyer, 1986), and then water can import the grain through pedicel xylem and phloem.

At 10 DAP, the soluble sugar content of JNK728 and XY779 grain was the maximum, and then decreased, while the starch content of grains began to increase rapidly at 10-15DAP (Fig. 9A, B). These findings further indicated that sugar began to convert promptly into starch and the filling rate was accelerated from grain formation (Fig. 10), which was identical with the results of previous studies (Huo, 2016; Li *et al*., 2016). Patrick and Offler (1995) found that when the polymer formation rate in pea cotyledons increased, so did the sucrose absorption rate of seed apoplasts. The osmotic pressure of apoplast sap decreases, which in turn stimulates the outflow of nutrients from seed coat to seed apoplast to match the rate of nutrient uptake by cotyledons (Patrick and Offler, 1995; Walker *et al*., 2000). In addition to providing nutrients, bulk flow of sap from the phloem also meets the water demand for volume growth in expansion sinks (Bret-Hart and Silk, 1994; Farrar *et al*., 1995; Pritchard *et al*., 2000). Studies had also shown that most of the water supplied to grain development might be transported through the phloem (Pate *et al*., 1977; Jenner, 1982). Therefore, it is conjectured that in the grain, when the transformation rate of sugar to starch is accelerated, the osmotic potential decreases, which promotes the transport of sugar in the P-C zone to the grain. This results in the decrease of the osmotic potential in the P-C zone, thus accelerates the import rate of the phloem. Then the water transported to the grain through the pedicel phloem is able to gradually meet the needs of grain development, and the pedicel xylem no longer transports water to the grain.

According to the results of Tang’s research (Tang, 2018), the grain rapid filling of sweet maize and common maize started synchronously. But the grain filling rate of sweet maize was lower than that of common maize. It may be deduced that the difference between the dye’s ability to move through the pedicel of sweet maize WT2000 at 20 DAP and the common maize JNK728 and XY779 only within 10 DAP may be due to the sweet maize’s lower rate of grain filling.The 100-grain dry weight of JNK728 and XY779 in 15 DAP was 7.23g and 6.80g respectively, the starch content was 40.57% and 38.56% respectively, and the 100-grain water content was 19.06g and 15.37g respectively. The 100-grain dry weight of sweet maize WT2000 in 20 DAP was 6.57g, the starch content was 36.64%, and the 100-grain water content was 20.36g. The accumulation of grain total dry matter and starch of sweet maize at 20 DAP was less than that of common maize at 15 DAP, while the grain water content of sweet maize was higher than that of common maize at 15 DAP. This also demonstrates the previous inference that in the early stage of grain development, the grain filling rate of sweet maize is lower than that of common maize. That is, the transport rate of pedicel phloem of sweet maize is lower, and pedicel xylem also continue to transport water to grain in order to meet grain development until the import of water through the pedicel phloem can satisfy the need of grain growth. In 20 DAP, the grain starch content of the ear middle position was higher than that of the ear bottom and apex position (Fig. 9C), that is to say, the grain filling rate of the ear middle position was higher than that of the ear bottom and apex position. The water entering grain of the ear middle position through the pedicel phloem is more, resulting in decline of that through the pedicel xylem, thus the dye color in pedicle xylem of the middle cob is lighter than that of the bottom and apex cob.

In this phase, the grain filling rate was not low because JNK728 grain was in the late stage of rapid filling and XY779 grain had finished rapid filling recently (Fig. 10). The grain volume was close to the maximum value (Fig. 8C), and the water content of grain and cob maximized in 20 DAP (Fig. 8A, B). The husk was not completely withered and loose, and the moisture content of husk was still above 50%(Fig. 12, Fig. 13). Less water is needed for ear development and lost by evaporation, and the water import through cob phloem meets or even exceeds the need of ear development in 25-30 DAP. Therefore, there is almost net water export from ear through the cob xylem, and dye movement can only be observed in the ear-pedicel or bottom of cob at this stage. When the grain drew to finish or had finished the rapid filling stage in 25-30 DAP, the soluble sugar content of grain began to increase, and reached the second peak for JNK728 and XY779 at 35 and 30DAP, respectively. The results show that from 25 DAP, the transformation rate of sugar to starch in grain begins to decrease, causing accumulation of soluble sugar and increase of soluble sugar content (Fig. 9). Then the content of soluble sugar in P-C region increases, resulting in the decrease of phloem unloading rate (Patrick and Offler, 1995; Lalonde *et al*., 2003; Bewley *et al*.,2013a; Patrick, 2013).

Lalonde *et al.* (2003) and Patrick (2013) considered that in the sink tissue, sucrose was unloaded from the phloem, reducing the solute concentration in the sieve tube, causing water diffusion from the sieve tube to the surrounding cells and reducing the pressure in the phloem. When the unloading rate of phloem decreases, the osmotic potential in the sieve tube decreases slowly, resulting in the decrease of the water absorption from xylem, thus the water import from plant to ear through cob xylem decreases or even stop. From pollination to grain formation, there is only net water import through cob xylem in the ear; from grain formation to 25 DAP, and from 30 DAP to physiological maturity, there is both water import and export through cob xylem in the ear; from 25 to 30DAP, there is only the net water export through cob xylem.

Some studies suggested that the reason of xylem transport decreasing could be the physical damage of xylem caused by growth strains (Lang and Düring, 1991; Coombe and McCarthy, 2000; Dražeta *et al*., 2004) or xylem blockage (Knipfer *et al*., 2015). Whereas, the “pressure flow hypothesis) (Crafts and Currier, 1963)held that the excess water was presumed to flow back to the parent plant through the xylem. The results of this study obviously support the “pressure flow hypothesis”, which indicates that water can not enter the ear through the cob xylem is not because the xylem is damaged or blocked, but the transport direction of water in the cob xylem has changed. The water loss of transpiration from the expanded leaves to the atmosphere begins to generate negative pressure (Px) (Jupa *et al*., 2016; Keller, 2020) in the xylem of the plant by hydraulic pressure, which promotes the surplus water in the apoplast of the sink tissue to flow back to the maternal plant through the xylem (Pate *et al*., 1985; Jenner and Jones, 1990; Keller, 2006; Zhang *et al*., 2006; Bewley *et al*., 2013a; Keller *et al*., 2015). The study of Zhang *et al.* (2007) suggested that the excess water in the sink tissue apoplast was transported to the phloem for recycle, which might involve the alternatiive opening and closing of aquaporins corresponding to phloem unloading and xylem circulation. Pate *et al.* (1985) showed the diurnal pattern of xylem cycle, which they attributed to changes in transpirational demand and phloem transport. Another possibility is that aquaporins are sensitive to osmotic potential in apoplast. Therefore, it is conjectured that in the process of grain filling, due to the rapid transformation of sugar to starch in the grain, the osmotic potential decreases, resulting in that the water in the grain returns to the P-C region, and then flows back to cob xylem driven by the negative pressure of the xylem produced by plant transpiration. Part of the water in the cob xylem is transported to the phloem through aquaporins and recycled, part flows back to the maternal plant through the xylem and is recycled by plant, and the remained is lost by evaporation.

The grain dehydration rate of JNK728 from 35 DAP to 45 DAP increased gradually, and that of XY790 from 30 DAP to 40 DAP remained stable at first, and then gradually increased (Fig. 10). The studies of Brooking (1990), Feng *et al*. (2017) and Gao *et al*. (2018) showed that grain dehydration was related to meteorological factors such as temperature and humidity. At the above stage, the moisture content of husk decreased gradually and began to loose rapidly, and the higher temperature and lower humidity in the dry shed promoted the grain dehydration, so the grain dehydration rate was accelerated in advance by surface evaporation. The grain coated experiment and the dye movement experiment were carried out in the field and in the dry shed, respectively. The temperature in the dry shed was higher than that in the field, which shortened the growth period of maize and led to the early end of the grain effective filling period and the early withering and loosening of husk. Therefore, the conversion time of the main driving force of grain dehydration is based on the results of grain coated experiment, that is, the mid-maturing cultivar JNK728 and the early-maturing cultivar XY790 begins to change at 5 and 10 days before physiological maturity, respectively.

Conclusion: when photosynthate is transported to the grain through the pedicel phloem, the osmotic pressure in the grain begins to increase, while the transport rate of the pedicel phloem is low, thus the pedicel xylem also transports water to the grain for its early expansion and development during the grain formation period (within about 10 DAP). With the end of the grain formation, the grain filling rate and the transport rate of pedicel phloem increase, bringing about the water transported to the grain through the pedicel xylem decreases gradually, and ceases at 15 DAP. When the grain filling rate was accelerated to that the water imported to grain through the pedicel phloem is capable to meet or even exceed the needs of grain development. Meanwhile the transformation rate of sugar to starch in grain is also accelerated, causing the decrease of osmotic potential in grain(Zhang *et al*., 2020) and the water backflow from grain to cob through the pedicel xylem. Some of the backflow water in cob is recycled by the phloem, some is lost through evaporation on the cob surface, and the rest flows back to the plant for reuse. With the advance of the grain filling process, the filling rate gradually decreases, and so dose the water requirement for grain development, thus the water imported through the pedicel phloem could still meet or even exceed the needs of grain development. Therefore, the surplus water in grain is continuously exported to the cob through the pedicel xylem, and this process continues until the physiological maturity. From the end of grain formation to physiological maturity, there is a very significant positive correlation between grain filling rate and dehydration rate. Combining with the results of grain coated experiment, it is proved that from grain formation to close to physiological maturity(5-10 days before physiological maturity), the main driving force of grain dehydration of the early and middle maturity maize cultivars is grain filling, and then changes to surface evaporation.

## Acknowledgements

We are grateful to the professors in the team of selection of high yield and suitable machine – harvesting maize cultivars for dense planting and related cultivation techniques making proposals for trials. We are also grateful to the Hebei Agricultural University which generously provided us with field sites for trials.

## Author contributions

F-LZ: funding acquisition; F-LZ: acquisition of the seed materials; F-LZ and G-PZ: designing the experiments; F-LZ, G-PZ, MM and W-WL: data collection; F-LZ and G-PZ: data analysis; G-PZ, F-LZ and MM: writing with contributions from all authors.

## Conflict of interest

The authors declare no conflict of interest.

## Funding

This work was supported by the National Key Research and Development Program of China (2016YFD0300300)

## Data availability

Data supporting the findings of this study are available from the corresponding author upon request.

## Notes

### Competing Interest Statement

The authors have declared no competing interest.

